# Tetraspanins from *Opisthorchis viverrini* stimulate cholangiocyte migration and inflammatory cytokine production

**DOI:** 10.1101/2023.06.12.544604

**Authors:** Apisit Ruangsuwast, Michael J. Smout, Paul J. Brindley, Alex Loukas, Thewarach Laha, Sujittra Chaiyadet

**Author notes:** **Address for correspondence:** Sujittra Chaiyadet, Department of Tropical Medicine, Faculty of Medicine, Khon Kaen University, Khon Kaen, 40002 Thailand.

## Abstract

The liver fluke *Opsithorchis viverrini* secretes extracellular vesicles (EVs) bearing CD63-like tetraspanins on their surface. Fluke EVs are actively internalized by host cholangiocytes in the bile ducts, where they drive pathology and promote neoplasia through induction of cellular proliferation and secretion of inflammatory cytokines. We investigated the effects of tetraspanins of the CD63 superfamily by co-culturing recombinant forms of the large extracellular loop (LEL) of *O. viverrini* tetraspanin-2 (rLEL-*Ov*-TSP-2) and tetraspanin-3 (rLEL-*Ov*-TSP-3) with non-cancerous human bile duct (H69) and cholangiocarcinoma (CCA, M213) cell lines. The results showed that cell lines co-cultured with excretory/secretory products from adult *O. viverrini* (*Ov-*ES) underwent significantly increased cell proliferation at 48 hours but not 24 hours compared to untreated control cells (*P*<0.05), whereas rLEL-*Ov*-TSP-3 co-culture resulted in significantly increased cell proliferation at both 24 hr (*P*<0.05) and 48 hr (*P*<0.01) time points. In like fashion, H69 cholangiocytes co-cultured with both *Ov*-ES and rLEL-*Ov*-TSP-3 underwent significantly elevated *Il-6* and *Il-8* gene expression for at least one of the time points assessed. Finally, both rLEL-*Ov*-TSP-and rLEL-*Ov*-TSP-3 significantly enhanced migration of both M213 and H69 cell lines. These findings indicated that *O. viverrini* CD63 family tetraspanins can promote a cancerous microenvironment by enhancing innate immune responses and migration of biliary epithelial cells.

---

Infection of humans with the liver fluke *Opisthorchis viverrini* is a major risk factor for the development of bile duct cancer, or cholangiocarcinoma (CCA) in fluke-endemic areas (Sripa et al. 2012, Brindley et al. 2021). The excretory/secretory (ES) products released by flukes when residing in the bile ducts have been shown to have carcinogenic properties (Sripa et al. 2007, Sripa et al. 2012, Chaiyadet et al. 2015a) that drive inflammation and induce proliferation of the biliary epithelial cells. *O. viverrini* ES products contain both soluble proteins and extracellular vesicles (EVs), the latter of which are released from the tegument and actively internalized by non-cancerous and cancerous human bile duct cell lines where they stimulate inflammatory cytokine secretion and abnormal cell growth (Chaiyadet et al. 2015b, Wilson and Jones 2021).

Tetraspanins are a highly conserved family proteins with four transmembrane domains, and two extracellular loops of unequal size: a small extracellular loop (SEL) and a large extracellular loop (LEL). The LEL region of TSPs contains between four and eight cysteine residues which from two to four disulfide bonds act as the “signal motif”. This allows for specific protein-protein interactions with adjacent proteins and other ligands (Levy and Shoham 2005). TSPs are involved in basic cell activities, including cell proliferation, cell fusion, motility, adhesion, migration, and signal transduction pathways (Hemler 2008). Previous studies have described three tetraspanins from the *O. viverrini* ES products belonging to the CD9 (*Ov*-TSP-1) (Piratae et al. 2012) and CD63 families (*Ov*-TSP-2 and *Ov*-TSP-3) (Chaiyadet et al. 2017). These TSPs are highly expressed in the parasite tegument throughout the liver fluke’s life cycle where they play essential roles in maintaining the integrity of the tegument and formation of membrane-bound EVs. Indeed, *Ov*-TSP-1 and TSP-2 have recently been identified as markers of EVs (Chaiyadet et al. 2022).

CD63 has been shown to be involved in metastasis of cancer cells (Seubert et al. 2015), where it acts as a pro-metastatic factor via □-catenin stabilization. While *O. viverrini* EVs are known to promote cell proliferation and inflammatory cytokine production in cholangiocytes, the specific role of CD63-like TSPs (which are abundant on the EV surface) in the development of *O. viverrini* infection-induced CCA has not been investigated. Herein we investigated the interactions between *O. viverrini* recombinant LEL domains of *Ov*-TSP-2 and TSP-3 and human normal cholangiocyte and CCA cell lines, and investigated their effects on cell migration and inflammatory cytokine production by cholangiocytes.

## MATERIALS AND METHODS

### Rodent model, parasite production and excretory/secretory products of *Opisthorchis viverrini*

Male Syrian golden hamsters 6-8 weeks were infected with 50 *O. viverrini* metacercariae by intragastric intubation (Sripa and Kaewkes 2000). Adult flukes were recovered from hamsters and cultured *in vitro* to produce ES products as described (Chaiyadet et al. 2019, Chaiyadet et al. 2022). The animals were housed and cared for in the animal facility at Faculty of Medicine, Khon Kaen University, in accordance with approved protocols from the Animal Ethics Committee of Khon Kaen University (IACUC-KKU-92/63).

### Recombinant protein production

The large extracellular loop of *O. viverrini* tetraspanin-2 (rLEL-*Ov*-TSP-2) and tetraspanin-3 (rLEL-*Ov*-TSP-3) were produced in recombinant form using *Pichia pastoris* as described (Phung et al. 2019) and *Escherichia coli* as described (Chaiyadet et al. 2017), respectively. In brief, LEL-*Ov-tsp-2* sequence was amplified by PCR and inserted into the pPICZαA plasmid. The recombinant plasmid was linearized with *Sac I* restriction enzyme after which it was used to transform *P. pastoris* X33 strain using electroporation (MicroPulser Electroporator, Bio-Rad, Hercules, CA, USA). Recombinant expression of transformed yeast was induced by 1% methanol for 5 days. Subsequently, the culture medium was clarified by centrifugation and then mixed with binding buffer (50 mM NaH2PO4, 300 mM NaCl, 5 mM Imidazole, pH 8.0) prior to chromatographic purification on Ni-NTA resin (Thermo Fisher Scientific, USA). After elution from the resin column, the purified protein was dialyzed against 20 mM HEPES using an Amicon ultra-15 (Merck, Rahway, NJ, USA).

The LEL-*Ov*-TSP-3 sequence was cloned into plasmid pET32a (Novagen, Madison, WI, USA) which was deployed to transform BL21DE32 strain *E. coli* for protein expression. The transformed bacteria were cultured in LB broth and induced to produce protein by the addition of IPTG (isopropylthio-β-galactoside) to 1 mM for 6 hours at 37 ºC. The bacterial pellet was harvested by centrifugation, recombinant proteins extracted from the cells, and recombinant LEL-*Ov*-TSP-3 isolated on Ni-NTA resin, under non-denaturing conditions using 500 mM imidazole for elution. The concentration of the purified protein was measured by NanoDrop 2000c spectrophotometer (Thermo Scientific, Waltham, MA, USA).

### Endotoxin measurement by limulus amebocyte lysate (LAL) assay

Recombinant proteins and *Ov*-ES products were evaluated for the presence of lipopolysaccharide (LPS) using a chromogenic LAL endotoxin assay kit (GenScript, Piscataway, NJ, USA) according to the manufacturer’s instructions. A standard curve was generated using LPS concentrations ranging from 0.1-0.01 endotoxin units (EU) per ml (0.01-0.001 ng/ml) for sample measurement. The absorbance of the reaction was measured at 545 nm using a microplate reader (Varioskan™ LUX microplate reader, Thermo Scientific).

### Cell proliferation assay

H69 cells were maintained in Dulbecco’s modified Eagle’s medium (DMEM)/Ham-F12 (Gibco, USA) supplemented with 10% fetal bovine serum (FBS), 100 Units/ml Penicillin and 100 Units/ml Streptomycin (Life Technologies, Carlsbad, CA, USA), insulin, adenine, epinephrine, T3-T, epidermal growth factors (EGF) and hydrocortisone (Ninlawan et al. 2010). The M213 CCA cell line was cultured in RPMI (Gibco, Thermo Fisher Scientific) supplemented with 10% FBS and 100 Units/ml Penicillin and 100 Units/ml Streptomycin.

Cell proliferation was measured using the MTT assay (Invitrogen, Thermo Fisher Scientific). Initially, 15,000 H69 cholangiocytes were seeded into 24 well plate and then grown overnight at 37 ºC in a 5% CO2 environment. Three hours prior to adding recombinant proteins or *Ov*-ES products, the growth media was replaced with a low nutrient of H69 media containing 0.5% FBS (Smout et al. 2015) for 3 hours before protein exposure. The cells were cultured with various concentrations (0.8-6.4 μg/ml) of rLEL-*Ov*-TSP-2, rLEL*Ov*-TSP-3, 1.25 μg/ml *Ov-*ES, or PBS in low nutrient media for 24 and 48 hours. Viable cells were quantified using the MTT assay with absorbance of 570 nm. The experiments were conducted in triplicate for each condition.

### Quantitative RT-PCR

H69 cholangiocytes were cultured with 1.6 μg/ml of the r*Ov*-TSP-2 and r*Ov*-TSP-3 in a low nutrient media for 24 or 48 hours then cells were preserved in Trizol reagent (Invitrogen, Life technologies) and stored at -80 °C until further processing. To measure cytokine expression, total cellular RNA was extracted using TriZol^®^ reagent and reverse transcribed to cDNA using a RevertAid First Strand cDNA Synthesis kit (Thermo Fisher Scientific). The cDNA was amplified using gene-specific primers designed to amplify a portion of the coding sequences. The amplification was performed using Maxima SyBr green qPCR master mix (Thermo Fisher Scientific). The qPCR was performed using a single 10 minute denaturation step at 95 °C, followed by 40 cycles of 30 sec at 94 °C, 30 sec at 55 °C, and 30 sec 72 °C, and a final extension step at 72 °C for 10 min. IL-6 primer sequence was: Forward; 5’-ACCCCTGACCCAACCACAAAT-3’, Reverse; 5’-CCTTAAAGCTGCGCAGAATGAGA- 3’; IL-8 primer sequence was Forward; 5’-GTGCAGTTTTGCCAAGGAGT-3’, Reverse; 5’-CTCTGCACCCAGTTTTCCTT-3’. Gene expression was normalized the □-actin gene, forward; 5’-TCCCTGGAGAAGAGCTACGA, Reverse; 5’AGCACTGTGTTGGCGTACAG.

### Transwell migration

The adenosquamous carcinoma cell line M213 which originated from human CCA tissue (Sripa et al. 2020) and H69 normal cholangiocytes were co-cultured with recombinant TSPs to assess cell migration. Briefly, 5×10^4^ of M213 or H69 cells were seeded in the upper chamber of a Transwell polycarbonate insert with 8.0 μm pore size (Corning, NY, USA). After 1 hour, rLEL-*Ov*-TSP-2, rLEL-*Ov*-TSP-3, or *Ov-*ES were added into to the lower chamber, and the cells were incubated for an additional 24 hours to allow migration to occur. The insert was removed, and cells on the upper side of the membrane were removed using a cotton bud. The membranes were fixed with 4% paraformaldehyde, washed with PBS, and stained with 5 μg/ml Hoechst for nuclear visualization. The migration of cells was assessed by counting the number of positive cells in ten random microscopy fields (10×) of each membrane. Images were taken using a Nikon Eclipse Ti microscope and analyzed with NIS-Elements version 4.3 software. The cells were counted using ImageJ version 1.52p.

### Statistical analysis

The data were presented as the mean ± standard error of three independent experiments and analyzed for normal distribution using GraphPad Prism using one-way ANOVA with post-hoc Tukey’s test. A *P* value of <0.05 was considered to be significant.

## RESULTS

### rLEL-*Ov*-TSP-3 and *Ov*-ES stimulates bile duct cell proliferation

The recombinant protein was assessed for endotoxin using LAL assay. The result showed that the LPS concentration in 1.6 μg/ml rLEL-*Ov*-TSP-2, 1.6 μg/ml rLEL-*Ov*-TSP-3 and 1.25 μg/ml *Ov-*ES was 0.019 ng/ml, 0.022 ng/ml, and 0.023 ng/ml respectively (Fig S1.). These values fall within the acceptable range for cell culture, as outlined by the manufacturer’s instructions.

To optimize the effect of protein concentration on H69 cell growth, rLEL-*Ov*-TSP-2 and rLEL-*Ov*-TSP-3 at various concentrations (0.8-6.4 μg/ml) were co-cultured with H69 cells and compared to the effect of ES products at a concentration of 1.25 μg/ml as described elsewhere (Chaiyadet et al. 2015a). In agreement with our earlier findings, *Ov-*ES also resulted in significantly elevated cell growth. Based on these findings, further studies were conducted using concentrations of 1.6 μg/ml of rLEL-*Ov*-TSP-2 and rLEL-*Ov*-TSP-3, and 1.25 μg/ml of *Ov*-ES in co-culture with cholangiocytes for 24 and 48 hours to investigate cell proliferation using MTT assay. rLEL-*Ov*-TSP-3 but not TSP-2 resulted in significantly enhanced cholangiocyte proliferation when compared to non-treated cells at 24 and 48 hours (Fig. 1).

**Fig. 1.**
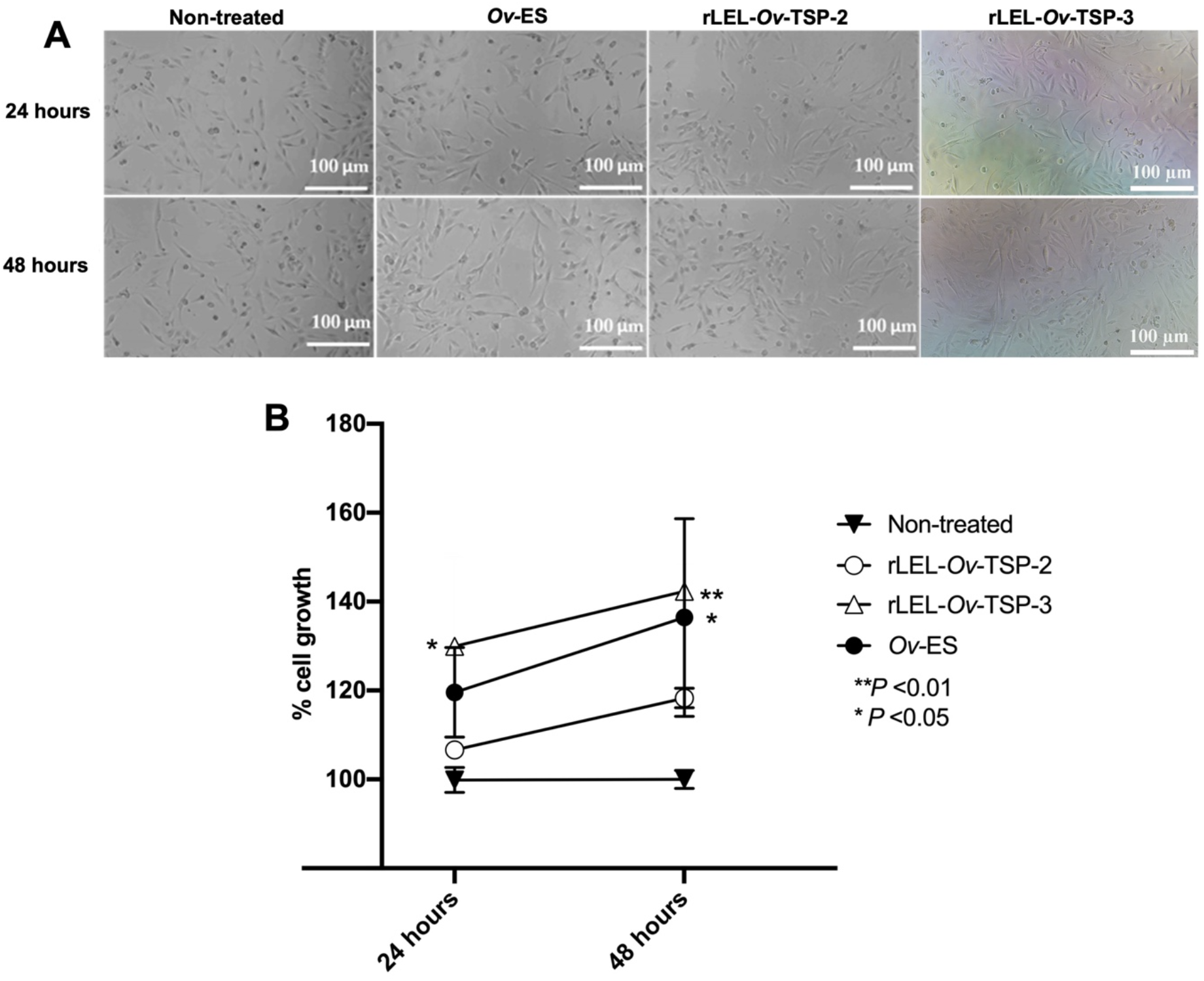
Growth of H69 cholangiocytes co-cultured with rLEL-*Ov*-TSP-2, rLEL-*Ov*-TSP-3 and *Ov-*ES using the MTT assay. The H69 cholangiocytes exposed with the 1.6 μg/ml rLEL-*Ov*-TSP-2 and rLEL-*Ov*-TSP-3, 1.25 μg/ml *Ov-*ES for 24 and 48 hours (A). The effect of rLEL-*Ov*-TSP-2, rLEL-*Ov*-TSP-3 on cholangiocyte growth compared to those exposed to *Ov-*ES as a positive control and non-treated as a negative control, using the MTT assay (B). \*\**P* <0.01, \**P* <0.05

### rLEL-*Ov*-TSP-3 and *Ov*-ES stimulate pro-inflammatory cytokine secretion by H69 cholangiocytes

To investigate the innate immune response of bile duct cells exposed to recombinant TSPs, gene expression levels of interleukin-6 (*Il-6*) and interleukin-8 (*Il-8*) were assessed after cholangiocytes were exposed to rLEL-*Ov*-TSP-2 and rLEL-*Ov*-TSP-3. Expression levels of the *Il-6* gene were significantly increased in H69 cells co-cultured with rLEL-*Ov*-TSP-3 after 24 hours and with *Ov*-ES after 48 hours, whereas rLEL-*Ov*-TSP-2 had no effect (Fig. 2A). Expression of the *Il-8* gene was significantly upregulated in H69 cells co-cultured with both rLEL-*Ov*-TSP-3 after 24 hours and *Ov*-ES after 24 and 48 hours. rLEL-*Ov*-TSP-2, however, did not induce changes in *Il-8* gene expression in cholangiocytes (Fig. 2B).

**Fig. 2.**
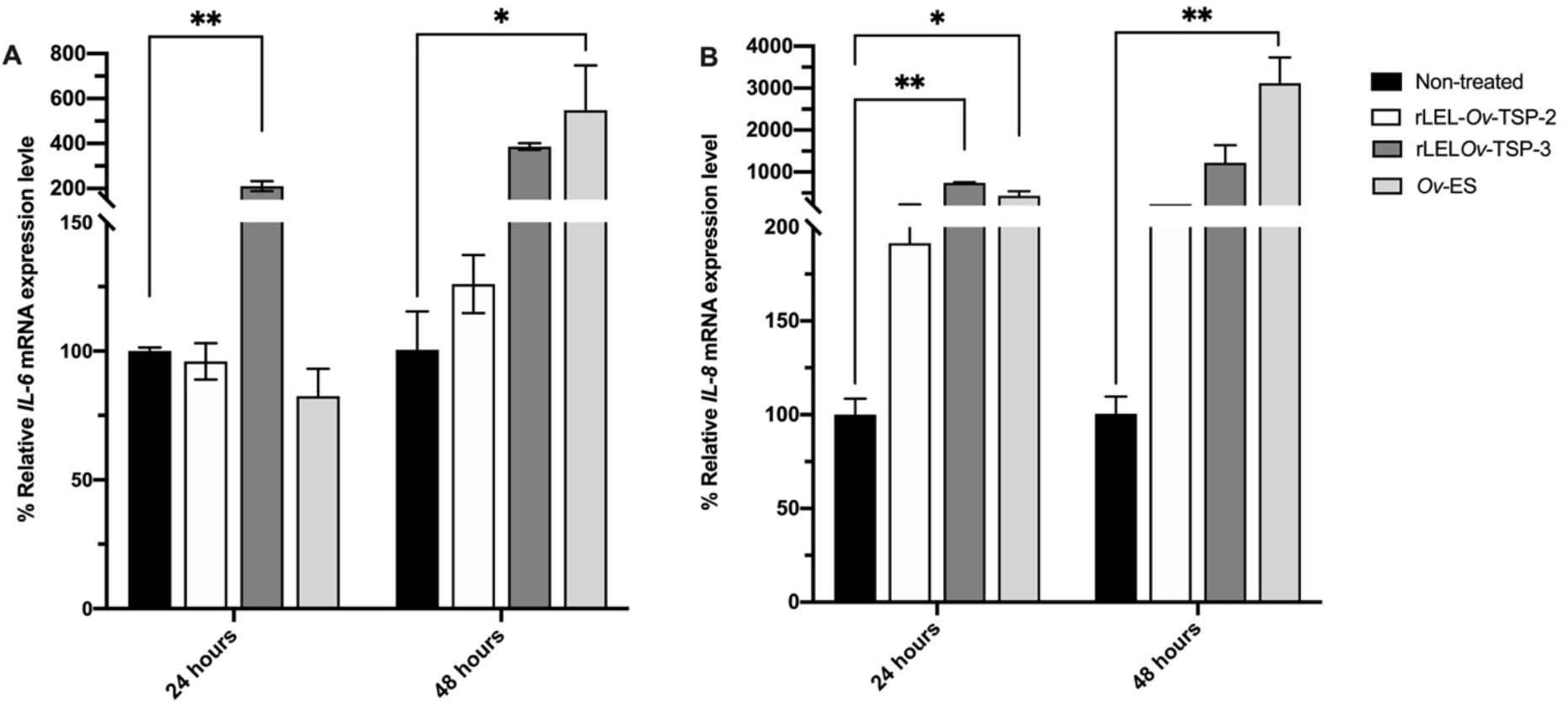
The mRNA expression levels of *Il-6* (A), and *Il-8* (B) in H69 cells after co-culture with rLEL-*Ov*-TSP-2, rLEL-*Ov*-TSP-3, and *O. viverrini* secretory products (*Ov*-ES) at 24 and 48 hours determined by qRT-PCR. \**P* < 0.05, \*\**P* < 0.01

### Tetraspanins induce cell migration in normal cholangiocyte and cholangiocarcinoma cell lines

To study the impact of *Ov-*TSPs on the migration of host cells, H69 cholangiocytes and M213 CCA cell lines were exposed to rLEL-*Ov*-TSP-2, rLEL-*Ov*-TSP-3, and *Ov*-ES for 24 hours using a Transwell migration assay (Techasen et al. 2014). The results indicated that both rLEL-*Ov*-TSP-2 and rLEL-*Ov*-TSP-3 induced significantly increased migration in H69 cholangiocyte (Fig. 3A) and M213 CCA (Fig. 3B) cell lines. However, *Ov*-ES only induced significant cell migration in M213 cells, but not in the cholangiocyte cell line.

**Fig. 3.**
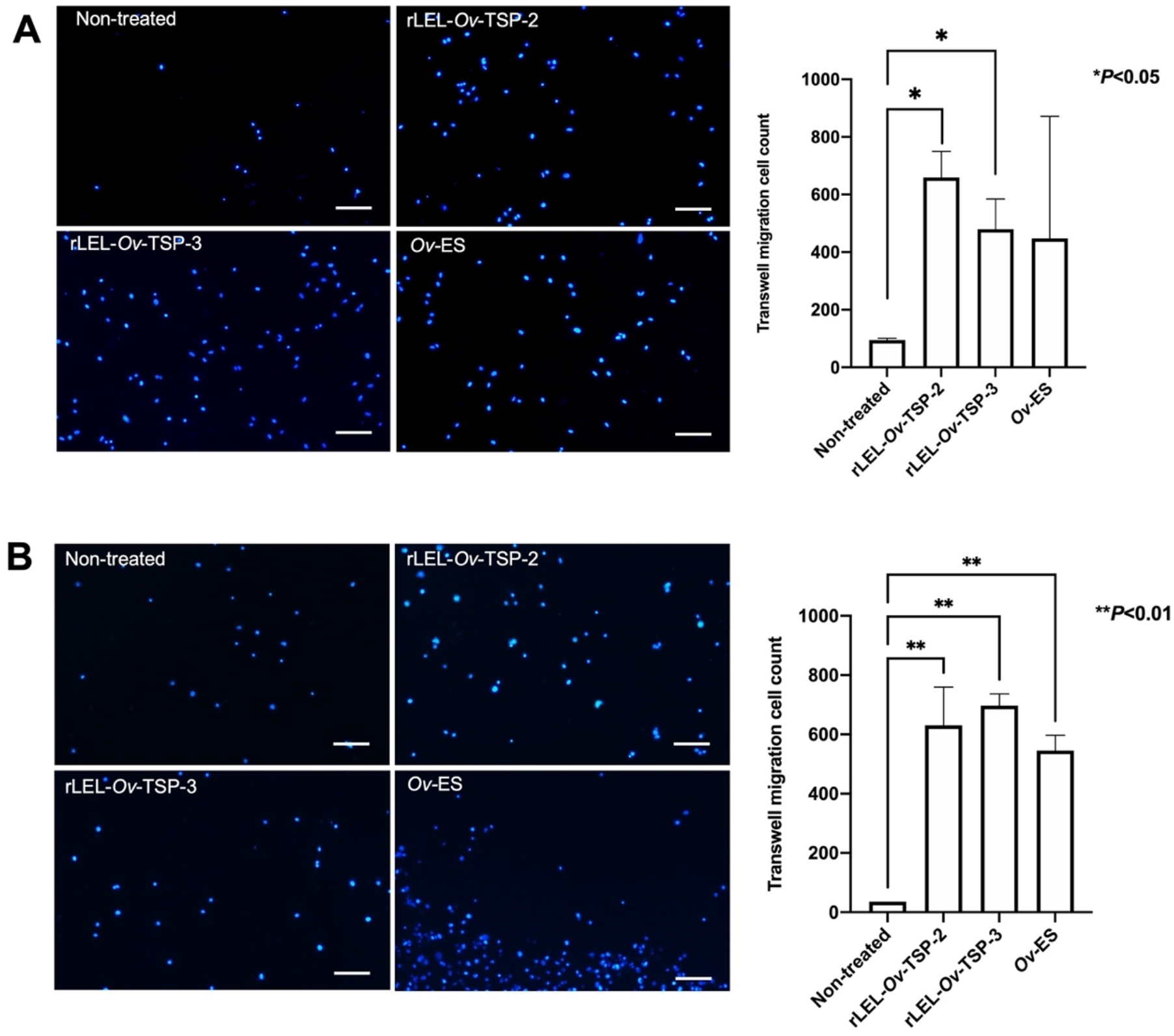
Tetraspanins of *O. viverrini* induced migration of normal and malignant cholangiocytes. The cholangiocyte H69 (A) and cholangiocarcinoma M213 (B) cell lines migrated through Transwell membranes following co-culture with rLEL-*Ov*-TSP-2, rLEL-*Ov*-TSP-3, and *Ov*-ES for 24 hours. Hoechst dye stained nuclei stained blue. ImageJ was used to count migrating cells \**P* < 0.05, \*\**P* <0.01.

## DISCUSSION

In helminths, TSPs play a key role in maintaining tegument biogenesis and stability (Chaiyadet et al. 2017) and are defined as markers of EVs (Chaiyadet et al. 2022). However, the effects of these proteins on cholangiocytes were, until now, unclear. Herein, two recombinant TSP LEL domains of *O. viverrini* were expressed and their impact on human cholangiocyte cell lines (normal and cancerous) were assessed.

Previous earlier reports revealed that *Ov*-ES stimulates proinflammatory cytokine secretion by cholangiocytes and induces abnormal cell growth (Ninlawan et al. 2010, Syal et al. 2012). Additionally, *Ov*-ES can increase the production of IL-6 and IL-8 from peripheral blood mononuclear cells of *O. viverrini*-infected human subjects (Surapaitoon et al. 2017). TSPs on the surface of *O. viverrini* extracellular vesicles (*Ov*-EVs) are important in the entry of vesicles into host target cells (Chaiyadet et al. 2022), whereupon the EVs trigger multiple pathways involved in cancer development, including Wnt signaling that can promote cell proliferation (Chaiyadet et al. 2015b). *Ov-*EVs that are taken up by cholangiocytes induce cell proliferation and secretion of IL-6 (Chaiyadet et al. 2015b), events that have been shown to promote tumorigenesis in liver fluke infection (Sripa et al. 2012). Antibodies against *Ov*-TSPs have been shown to bind to the EV surface and block EV internalization by cholangiocytes, thereby attenuating cell proliferation and reducing IL-6 production (Chaiyadet et al. 2015b, Chaiyadet et al. 2022).

The interactions of TSPs with various binding partners, including integrins and other TSPs triggers cellular activity such as downstream signaling in response to migratory signals (Hemler 2008, Bassani and Cingolani 2012, Schroder et al. 2013). This regulation occurs in both normal and pathological processes such as cancer metastasis and inflammation. For example, CD151 promotes migration of epidermoid carcinoma cells via α3β1 andα6β4 integrin-dependent cell adhesion and migration (Hong et al. 2012). A number of studies have shown that N-glycosylation in LELs of TSPs regulates adhesion and motility of the cell by binding with α3 and α5 integrin domains (Ono et al. 2000). Moreover, the interaction of CD63 to CXCR4 via the N-glycans triggers downstream signaling of the chemokine receptor (Yoshida et al. 2009). The large extracellular loop of most TSPs are glycosylated via one or more potential N-linked glycosylation site (Wang et al. 2012, Marjon et al. 2016), and *O. viverrini* TSP LELs are also predicted to be glycosylated via the presence of asparagine residues (Chaiyadet et al. 2015a).

*Ov*-TSP-3 induced upregulated expression in cholangiocytes of *Il-8*, a potent chemoattractant, implying that this process may drive cell migration through the chemokine receptors, CXCR1 and CXCR2, both of which are highly expressed in cancer cells and can trigger liver cell migration and invasion (Bi et al. 2019). Taken together, *O. viverrini* TSPs may interact with their partner molecules in cholangiocytes and stimulate downstream signals that promote cell migration, and ultimately contribute to malignant transformation.

In conclusion, *Ov*-TSP-2 and *Ov*-TSP-3 stimulate cell proliferation and increase the production of the pro-inflammatory cytokines, IL-6 and IL-8, leading to increased migration of both normal cholangiocyte and cholangiocarcinoma cell lines. These processes contribute to the known carcinogenic properties of liver fluke infection and help to explain why this group of parasites is recognized as a group 1 biological carcinogen (Humans 2012). Our findings also highlight the importance of interrupting the molecular interactions between fluke EVs and host biliary cells by vaccination (Chaiyadet et al. 2019, Phung et al. 2019, Phumrattanaprapin et al. 2021a, Phumrattanaprapin et al. 2021b) to develop therapeutic strategies that prevent this carcinogenic infection.

## Acknowledgments

We express our gratitude to the laboratory staff of the Department of Parasitology and Department of Tropical Medicine, Faculty of Medicine, Khon Kaen University for their facilitation and technical support.

## Funding

This study was supported by the National Cancer Institute, National Institutes of Health, award R01 CA164719 (TL, AL and PJB) and Research and Graduate Studies, Khon Kaen University (TL and SC).

## Author Contributions

A.P., S.C., and T.L. conceived the project and designed the experiments. A.P. and S.C. performed the experiments and analyzed the results. All authors reviewed the manuscript.

## Figure legends

**Fig. S1.**
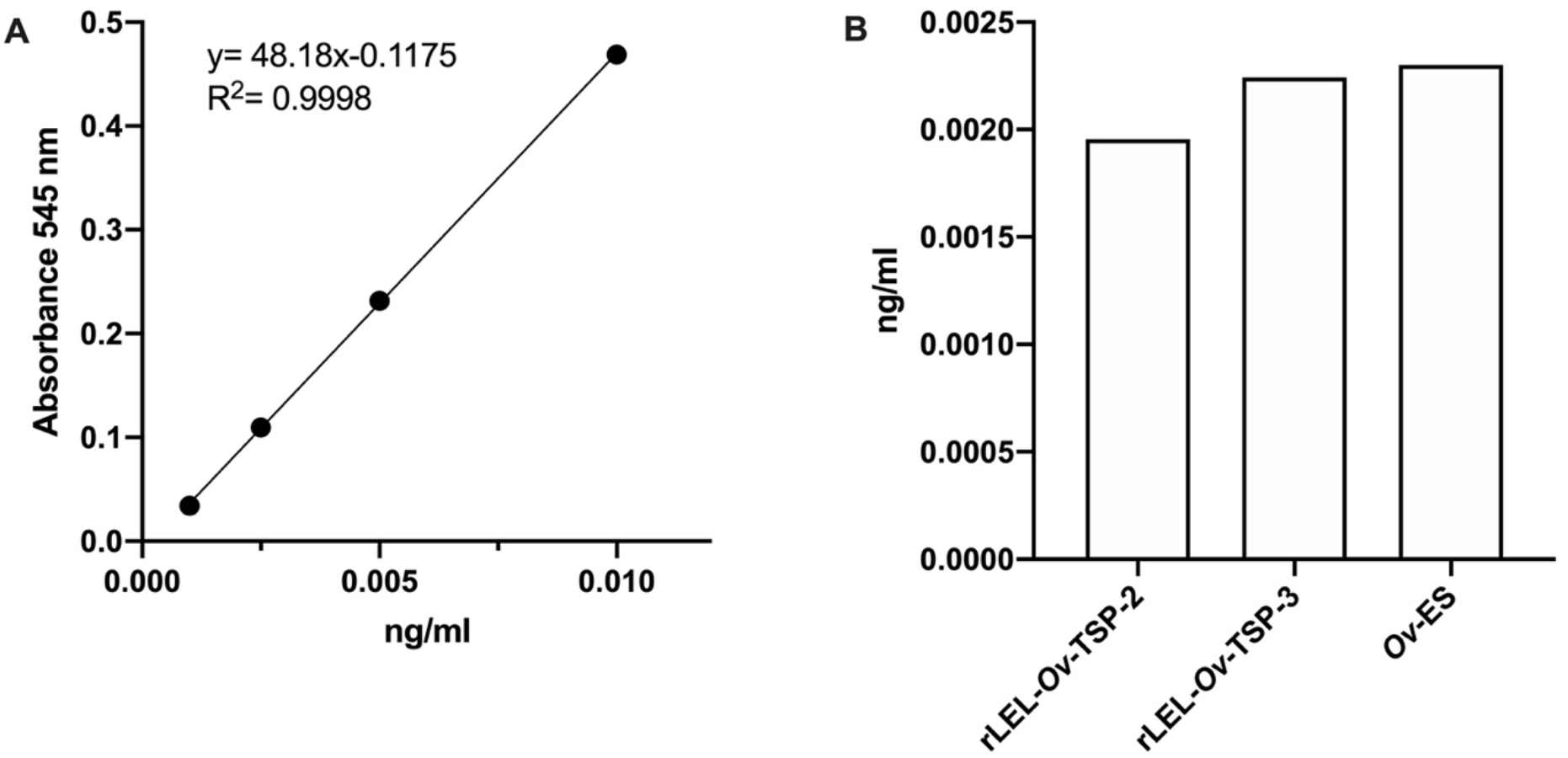
A standard curve of endotoxin concentration at 0.01-0.001 ng/ml was developed using the Limulus Amebocyte lysate (LAL) assay (A) to measure the endotoxin levels of rLEL-*Ov*-TSP2, rLEL-*Ov*-TSP-3, and *Ov*-ES (B).

## Notes

### Competing Interest Statement

The authors have declared no competing interest.

## REFERENCES

Bassani S., Cingolani L.A. 2012: Tetraspanins: Interactions and interplay with integrins. Int J Biochem Cell Biol 44: 703–708.

Bi H., Zhang Y., Wang S., Fang W., He W., Yin L., Xue Y., Cheng Z., Yang M., Shen J. 2019: Interleukin-8 promotes cell migration via CXCR1 and CXCR2 in liver cancer. Oncol Lett 18: 4176–4184.

Brindley P.J., Bachini M., Ilyas S.I., Khan S.A., Loukas A., Sirica A.E., Teh B.T., Wongkham S., Gores G.J. 2021: Cholangiocarcinoma. Nat Rev Dis Primers 7: 65.

Chaiyadet S., Krueajampa W., Hipkaeo W., Plosan Y., Piratae S., Sotillo J., Smout M., Sripa B., Brindley P.J., Loukas A., Laha T. 2017: Suppression of mRNAs encoding CD63family tetraspanins from the carcinogenic liver fluke Opisthorchis viverrini results in distinct tegument phenotypes. Sci Rep 7: 14342.

Chaiyadet S., Smout M., Johnson M., Whitchurch C., Turnbull L., Kaewkes S., Sotillo J., Loukas A., Sripa B. 2015a: Excretory/secretory products of the carcinogenic liver fluke are endocytosed by human cholangiocytes and drive cell proliferation and IL6 production. Int J Parasitol 45: 773–781.

Chaiyadet S., Sotillo J., Krueajampa W., Thongsen S., Brindley P.J., Sripa B., Loukas A., Laha T. 2019: Vaccination of hamsters with Opisthorchis viverrini extracellular vesicles and vesicle-derived recombinant tetraspanins induces antibodies that block vesicle uptake by cholangiocytes and reduce parasite burden after challenge infection. PLoS Negl Trop Dis 13: e0007450.

Chaiyadet S., Sotillo J., Krueajampa W., Thongsen S., Smout M., Brindley P.J., Laha T., Loukas A. 2022: Silencing of Opisthorchis viverrini Tetraspanin Gene Expression Results in Reduced Secretion of Extracellular Vesicles. Front Cell Infect Microbiol 12: 827521.

Chaiyadet S., Sotillo J., Smout M., Cantacessi C., Jones M.K., Johnson M.S., Turnbull L., Whitchurch C.B., Potriquet J., Laohaviroj M., Mulvenna J., Brindley P.J., Bethony J.M., Laha T., Sripa B., Loukas A. 2015b: Carcinogenic Liver Fluke Secretes Extracellular Vesicles That Promote Cholangiocytes to Adopt a Tumorigenic Phenotype. J Infect Dis 212: 1636–1645.

Hemler M.E. 2008: Targeting of tetraspanin proteins--potential benefits and strategies. Nat Rev Drug Discov 7: 747–758.

Hong I.K., Jeoung D.I., Ha K.S., Kim Y.M., Lee H. 2012: Tetraspanin CD151 stimulates adhesion-dependent activation of Ras, Rac, and Cdc42 by facilitating molecular association between beta1 integrins and small GTPases. J Biol Chem 287: 32027–32039.

Humans I.W.G.o.t.E.o.C.R.t. 2012: Biological agents. IARC Monogr Eval Carcinog Risks Hum 100: 1–441.

Levy S., Shoham T. 2005: The tetraspanin web modulates immune-signalling complexes. Nat Rev Immunol 5: 136–148.

Marjon K.D., Termini C.M., Karlen K.L., Saito-Reis C., Soria C.E., Lidke K.A., Gillette J.M. 2016: Tetraspanin CD82 regulates bone marrow homing of acute myeloid leukemia by modulating the molecular organization of N-cadherin. Oncogene 35: 4132–4140.

Ninlawan K., O’Hara S.P., Splinter P.L., Yongvanit P., Kaewkes S., Surapaitoon A., LaRusso N.F., Sripa B. 2010: Opisthorchis viverrini excretory/secretory products induce tolllike receptor 4 upregulation and production of interleukin 6 and 8 in cholangiocyte. aParasitol Int 59: 616–621.

Ono M., Handa K., Withers D.A., Hakomori S. 2000: Glycosylation effect on membrane domain (GEM) involved in cell adhesion and motility: a preliminary note on functional alpha3, alpha5-CD82 glycosylation complex in ldlD 14 cells. Biochem Biophys Res Commun 279: 744–750.

Phumrattanaprapin W., Chaiyadet S., Brindley P.J., Pearson M., Smout M.J., Loukas A., Laha T. 2021a: Orally Administered Bacillus Spores Expressing an Extracellular Vesicle-Derived Tetraspanin Protect Hamsters Against Challenge Infection With Carcinogenic Human Liver Fluke. J Infect Dis 223: 1445–1455.

Phumrattanaprapin W., Pearson M., Pickering D., Tedla B., Smout M., Chaiyadet S., Brindley P.J., Loukas A., Laha T. 2021b: Monoclonal Antibodies Targeting an Opisthorchis viverrini Extracellular Vesicle Tetraspanin Protect Hamsters against Challenge Infection. Vaccines (Basel) 9.

Phung L.T., Chaiyadet S., Hongsrichan N., Sotillo J., Dieu H.D.T., Tran C.Q., Brindley P.J., Loukas A., Laha T. 2019: Recombinant Opisthorchis viverrini tetraspanin expressed in Pichia pastoris as a potential vaccine candidate for opisthorchiasis. Parasitol Res 118: 3419–3427.

Piratae S., Tesana S., Jones M.K., Brindley P.J., Loukas A., Lovas E., Eursitthichai V., Sripa B., Thanasuwan S., Laha T. 2012: Molecular characterization of a tetraspanin from the human liver fluke, Opisthorchis viverrini. PLoS Negl Trop Dis 6: e1939.

Schroder H.M., Hoffmann S.C., Hecker M., Korff T., Ludwig T. 2013: The tetraspanin network modulates MT1-MMP cell surface trafficking. Int J Biochem Cell Biol 45: 1133–1144.

Seubert B., Cui H., Simonavicius N., Honert K., Schafer S., Reuning U., Heikenwalder M., Mari B., Kruger A. 2015: Tetraspanin CD63 acts as a pro-metastatic factor via betacatenin stabilization. Int J Cancer 136: 2304–2315.

Smout M.J., Sotillo J., Laha T., Papatpremsiri A., Rinaldi G., Pimenta R.N., Chan L.Y., Johnson M.S., Turnbull L., Whitchurch C.B., Giacomin P.R., Moran C.S., Golledge J., Daly N., Sripa B., Mulvenna J.P., Brindley P.J., Loukas A. 2015: Carcinogenic Parasite Secretes Growth Factor That Accelerates Wound Healing and Potentially Promotes Neoplasia. PLoS Pathog 11: e1005209.

Sripa B., Brindley P.J., Mulvenna J., Laha T., Smout M.J., Mairiang E., Bethony J.M., Loukas A. 2012: The tumorigenic liver fluke Opisthorchis viverrini--multiple pathways to cancer. Trends Parasitol 28: 395–407.

Sripa B., Kaewkes S. 2000: Localisation of parasite antigens and inflammatory responses in experimental opisthorchiasis. Int J Parasitol 30: 735–740.

Sripa B., Kaewkes S., Sithithaworn P., Mairiang E., Laha T., Smout M., Pairojkul C., Bhudhisawasdi V., Tesana S., Thinkamrop B., Bethony J.M., Loukas A., Brindley P.J. 2007: Liver fluke induces cholangiocarcinoma. PLoS Med 4: e201.

Sripa B., Seubwai W., Vaeteewoottacharn K., Sawanyawisuth K., Silsirivanit A., Kaewkong W., Muisuk K., Dana P., Phoomak C., Lert-Itthiporn W., Luvira V., Pairojkul C., Teh B.T., Wongkham S., Okada S., Chamgramol Y. 2020: Functional and genetic characterization of three cell lines derived from a single tumor of an Opisthorchis viverrini-associated cholangiocarcinoma patient. Hum Cell 33: 695–708.

Surapaitoon A., Suttiprapa S., Khuntikeo N., Pairojkul C., Sripa B. 2017: Cytokine profiles in Opisthorchis viverrini stimulated peripheral blood mononuclear cells from cholangiocarcinoma patients. Parasitol Int 66: 889–892.

Syal G., Fausther M., Dranoff J.A. 2012: Advances in cholangiocyte immunobiology. Am J Physiol Gastrointest Liver Physiol 303: G1077–1086.

Techasen A., Loilome W., Namwat N., Khuntikeo N., Puapairoj A., Jearanaikoon P., Saya H., Yongvanit P. 2014: Loss of E-cadherin promotes migration and invasion of cholangiocarcinoma cells and serves as a potential marker of metastasis. Tumour Biol 35: 8645–8652.

Wang H., Zhang W., Zhao J., Zhang L., Liu M., Yan G., Yao J., Yu H., Yang P. 2012: N-Glycosylation pattern of recombinant human CD82 (KAI1), a tumor-associated membrane protein. J Proteomics 75: 1375–1385.

Wilson R.A., Jones M.K. 2021: Fifty years of the schistosome tegument: discoveries, controversies, and outstanding questions. Int J Parasitol 51: 1213–1232.

Yoshida T., Ebina H., Koyanagi Y. 2009: N-linked glycan-dependent interaction of CD63 with CXCR4 at the Golgi apparatus induces downregulation of CXCR4. Microbiol Immunol 53: 629–635.

